# Reactive oxygen species as important regulators of cell division

**DOI:** 10.1101/2020.03.06.980474

**Authors:** Weiliang Qi, Li Ma, Fei Wang, Ping Wang, Junyan Wu, Jiaojiao Jin, Songqing Liu, Wancang Sun

**Author notes:** Gansu Agricultural University, Anning District of Lanzhou, China. Corresponding Author: Dr. Wancang Sun.

## Abstract

Currently, the role of reactive oxygen species (ROS) in plant growth is a topic of interest. In this study, we discuss the role of ROS in cell division. We analyzed ROS’ impact on the stiffness of plant cell walls and whether ROS play an important role in *Brassica napus*’ ability to adapt to cold stress. Cultivated sterile seedlings and calli of cold-tolerant cultivar 16NTS309 were subjected to cold stress at 25°C and 4°C, respectively. Under normal conditions, O^2.−^ mainly accumulated in the leaf edges, shoot apical meristem, leaf primordia, root tips, lateral root primordia, calli of meristematic nodular tissues, cambia, vascular bundles and root primordia, which are characterized by high division rates. After exposure to cold stress, the malondialdehyde and ROS (O^2.−^) contents in roots, stems and leaves of cultivar 16NTS309 were significantly higher than under non-cold conditions (*P* < 0.05). ROS (O^2.−^) were not only distributed in these zones, but also in other cells, at higher levels than under normal conditions. A strong ROS-based staining appeared in the cell wall. The results support a dual role for apoplastic ROS, in which they have direct effects on the stiffness of the cell wall, because ROS cleave cell-wall, and act as wall loosening agents, thereby either promoting or restricting cellular division. This promotes the appearance of new shoots and a strong root system, allowing plants to adapt to cold stress.

---

All living organisms, including plants, face extremes of sudden and adverse environmental conditions, such as cold, but unlike animals, plants are sessile and, therefore, cannot move to avoid stress [1, 2]. Cold stress is an abiotic stress that severely negatively impacts the growth and development of plants during any developmental stage, germination, seedling, vegetative, reproductive or grain maturity, leading to a reduction in grain yield. However, cold stress is one of a multiplex of factors in a specific environment that affects plants [3] and results in different types of reactive oxygen species (ROS) production in plants, including singlet oxygen, superoxide anion (O^2.−^), hydrogen peroxide (H_2_O_2_) and hydroxyl radicals [4-7]. Currently, the role of reactive oxygen species (ROS) in plant growth is a topic of interest. ROS not only act as signals in cells that activate a number of different defense mechanisms to protect cells[8], but also damage cells. ROS signaling mechanisms also play important roles in regulating the balance between cell proliferation and differentiation in both animals and plants [9]. In *Drosophila*, changing ROS levels can switch the status of hematopoietic cells from proliferation to differentiation[10]. ROS have direct effects on the stiffness of the plant cell wall, either promoting or restricting cellular division [11, 12]. Thus, it is necessary to determine and understand any correlation between ROS and cell division.

Winter rapeseed (*Brassica napus*), as a cover crop, helps to eliminate a dust source for the damaging sand storms in northern China. It is not only possible, but also beneficial economically, environmentally and ecologically, to grow winter rapeseed in dry and cold regions in northwestern China [13, 14]. However, the low temperatures in the winter make it difficult for *B. napus* varieties to survive [7, 13]. To improve the plant’s resistance to cold stress, the antioxidant capacity needs to be increased as does our understanding of the role of ROS in plant growth. Tissue culture-based approaches are convenient to operate under controlled environmental conditions, requiring less time and space (Fig 1). They are used to investigate the physiology and biochemistry of plants cultured under various environmental stress conditions [15]. To ensure the veracity of the experiment data, sterile seedlings need to be cultivated, unlike in field trials [16]. The advantage of this method is that ROS production, in response to biotic or abiotic stresses that occur when materials are treated in the natural environment, can be excluded.

**Fig 1.**
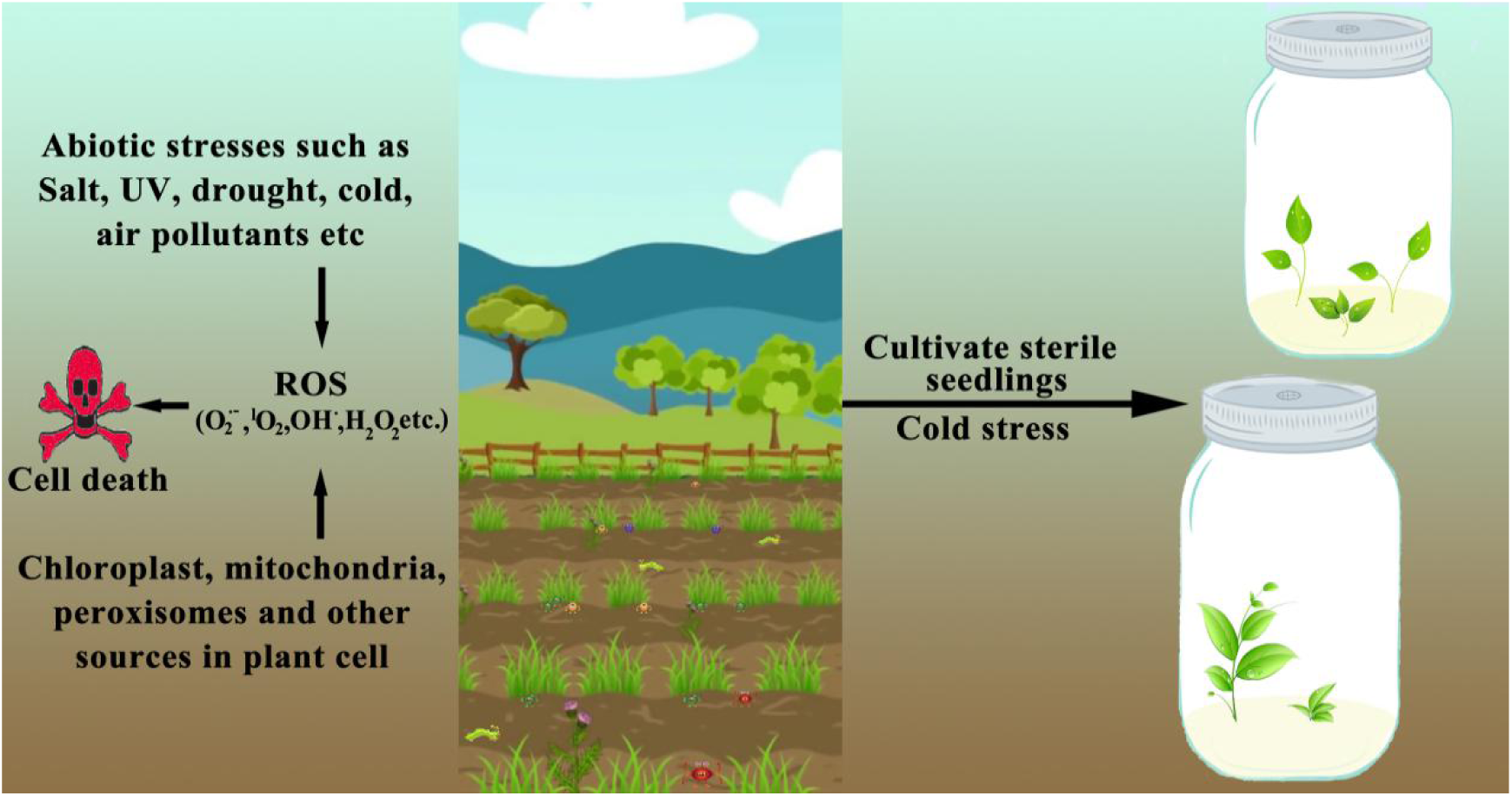
Cultivating sterile seedlings and calli of *B. napus* cultivar 16NTS309. Abiotic and biotic stresses are factors in a specific environment that can severely negatively affect plant growth, resulting in the production of different types of ROS. Tissue culture approaches have proven to be more convenient under controlled environmental conditions, requiring limited time and space. The advantage of this method is that it can exclude ROS production in response to biotic stresses that occur when materials are treated in a natural environment.

Our research group studied the feasibility of expanding winter rapeseed northwards into cold regions in northwestern China and bred new lines of *B. napus* having a strong cold tolerance, which could over winter in the 36°03′N area at an altitude of 2,150 m. These are the essential *B. napus* germplasm resources having strong cold tolerance levels used for breeding in northern China [17]. Physiological responses to low temperatures in *B. napus* have been intensively investigated, but the ROS signaling mechanisms underlying cold tolerance and resistance in plants is still rather poorly understood. In this study, we address several key questions: What is the correlation between ROS and cell division? Do ROS signaling mechanisms play important roles in the stiffness of the plant cell walls, thereby promoting root and leaf growth and differentiation? Does this promote new shoots and strong root systems that are required to adapt to cold stress? The aim was to provide new insights on the physiological and biochemical mechanisms and cytology associated with responses of *B. napus* to cold stress for use in breeding cold-resistant varieties.

## 1 Materials and methods

### 1.1 Cultivating sterile seedlings and calli of 16NTS309

The *B. napus* cultivar 16NTS309 (strongly resistant to cold damage) was produced by the Key Laboratory of Crop Genetics Improvement and Germplasm Enhancement of Gansu Province, Lanzhou. Plant sterile seedlings and calli of *B. napus* cultivar 16NTS309 were established using seeds and leaves. Initially, to produce sterile seedlings, seeds were soaked for 8 h and surface-sterilized with 75% alcohol for 30 s, followed by 0.5% HClO for 8 min. Then, the seeds were placed in 200-mL flasks containing 50 mL of liquid MS medium. The medium containing MS salts was supplemented with 15 g·L^−1^ saccharose and 17.5 g·L^−1^ agar, adjusted to pH 6.0 with NaOH or HCl and autoclaved at 121°C for 20 min. The growth conditions were 18-h light: 6-h dark photocycles at 22°C. In total, 100 sterile seedlings were obtained after 5 d of incubation. Then, calli were induced from leaves of sterile seedlings that were placed on callus-generating medium containing MS salts supplemented with 1 mg·L^−1^ 2,4-dichlorophenoxyacetic acid (2,4-D)and 1 mg·L^−1^6-benzylaminopurine (6-BA) that had been adjusted to pH 6.0 with NaOH or HCl and then autoclaved at 121°C for 20 min. In total, 90 calli were obtained after 7–14 d of incubation. Finally, sterile seedlings and calli of *B. napus* cultivar16NTS309 were divided into two groups: one was maintained at 25°C as the non-cold control and the other was subjected to cold stress at 4°C. To determine whether ROS were produced in the extracellular and intracellular environment, we conducted the same experiments using onions and subjected them to histochemical staining.

### 1.2 Physiological index method

Indexes of the malondialdehyde (MDA) and ROS (O^2.−^) contents were analyzed as indicators of physiological responses[18, 19]. Root, stem and leaf samples from cultivated sterile 16NTS309 seedlings were taken for morphological and physiological these analyses. The indexes were statistically analyzed using SPSS 19.0.

### 1.3 Detection of ROS (O^2.−^)

To determine the distribution of ROS (O^2.−^), in the plants, we stained plants with nitroblue tetrazolium (NBT), which is widely used as an indicator of ROS (O^2.−^) levels. The detection of ROS (O^2.−^) was assessed as previously described [20]. Briefly, plants were immersed for 8 h in 1 mg·mL^−1^ NBT staining solution, which was protected from light. After infiltration, the stained plants were bleached in an acetic acid:glycerol:ethanol (1:1:3, v/v/v) solution at 100°C for 10–20 min and then stored in 95% (v/v) ethanol until scanned.

### 1.4 Tissue section

NBT specifically reacts with O^2.−^ and forms a blue formazan precipitate. The 16NTS309 root, leaf and callus samples having the deepest blue formazan precipitate were sectioned for the morphoanatomical analysis. These samples were fixed in 50% mixed liquor of formalin–aceticacid–alcohol (FAA), softened using 20% ethylenediamine and embedded in paraffin. The samples were sectioned using a rotary microtome and stained with PAS, naphthol yellow or saffron, solid green staining. Slides were mounted in synthetic resin and images were captured using a digital image acquisition system.

## 2. Results

### 2.1 The physiological indexes of MDA and ROS (O^2.−^)

Under cold-stress conditions, the physiological indexes of MDA and ROS (O^2.−^) in roots, stems and leaves of cultivar 16NTS309 were measured. After 48 h of cold stress, the MDA and ROS (O^2.−^) contents were significantly higher in roots, stems and leaves compared with under non-cold conditions (*P* < 0.05). Under cold-stress conditions, the ROS contents in 16NTS309 roots, stems and leaves reached 216.397 µg·g^−1^ fresh weight (FW), 163.267 µg·g^−1^ FW and 217.627 µg·g^−1^ FW, respectively (Fig 2A). The MDA contents in roots and leaves of 16NTS309 reached 0.61 µmol·g^−1^ and 0.55 µmol·g^−1^, respectively, which were higher than the level in stems (Fig 2B).

**Fig 2.**
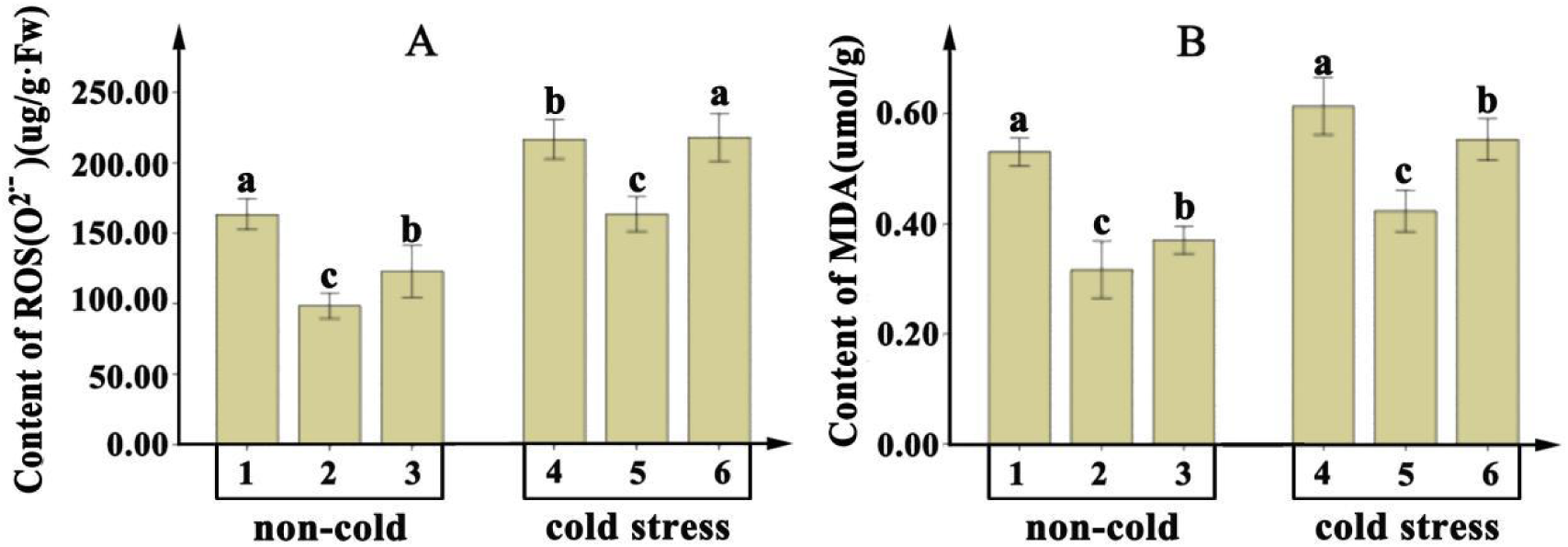
The physiological indexes of MDA and ROS (O^2.−^) were analyzed in root (1 and 4), stem (2 and 5) and leaf (3 and 6) samples from cultivated sterile *Brassica napus* 16NTS309 seedlings exposed to non-cold and cold-stress conditions.

### 2.2 Detection of ROS (O^2.−^) in leaves and roots

The shoot apical meristem (SAM) is the distal-most portion of the shoot and comprises two groups of cells: the initial or source cells and the progenitor cells of tissues and lateral organs [21, 22] By contrast, the shoot apex comprises several cell and tissue types: the SAM itself, a region just proximal to the meristem, in which lateral organ primordia, such as leaf primordia (LPs), form, a sub-apical region, in which the shoot widens and primordia enlarge, and the region of maturation, in which differentiation becomes apparent [21, 22]. We found a similar phenomenon in *B. napus* cultivar 16NTS309. Under normal conditions, the main ROS (O^2.−^) concentrations were in leaf edges (Fig 3a, e, f), SAM and LPs (Fig 3c, d). This organization was the same as cells having a strong division capacity. Presumably there is a correlation between ROS (O^2.−^) and cell division. After exposure to cold stress, the ROS (O^2.−^) level increased in leaves (Fig 3g, k, l), LPs and SAM (Fig 3 i, j) to greater levels than under normal conditions.

**Fig 3.**
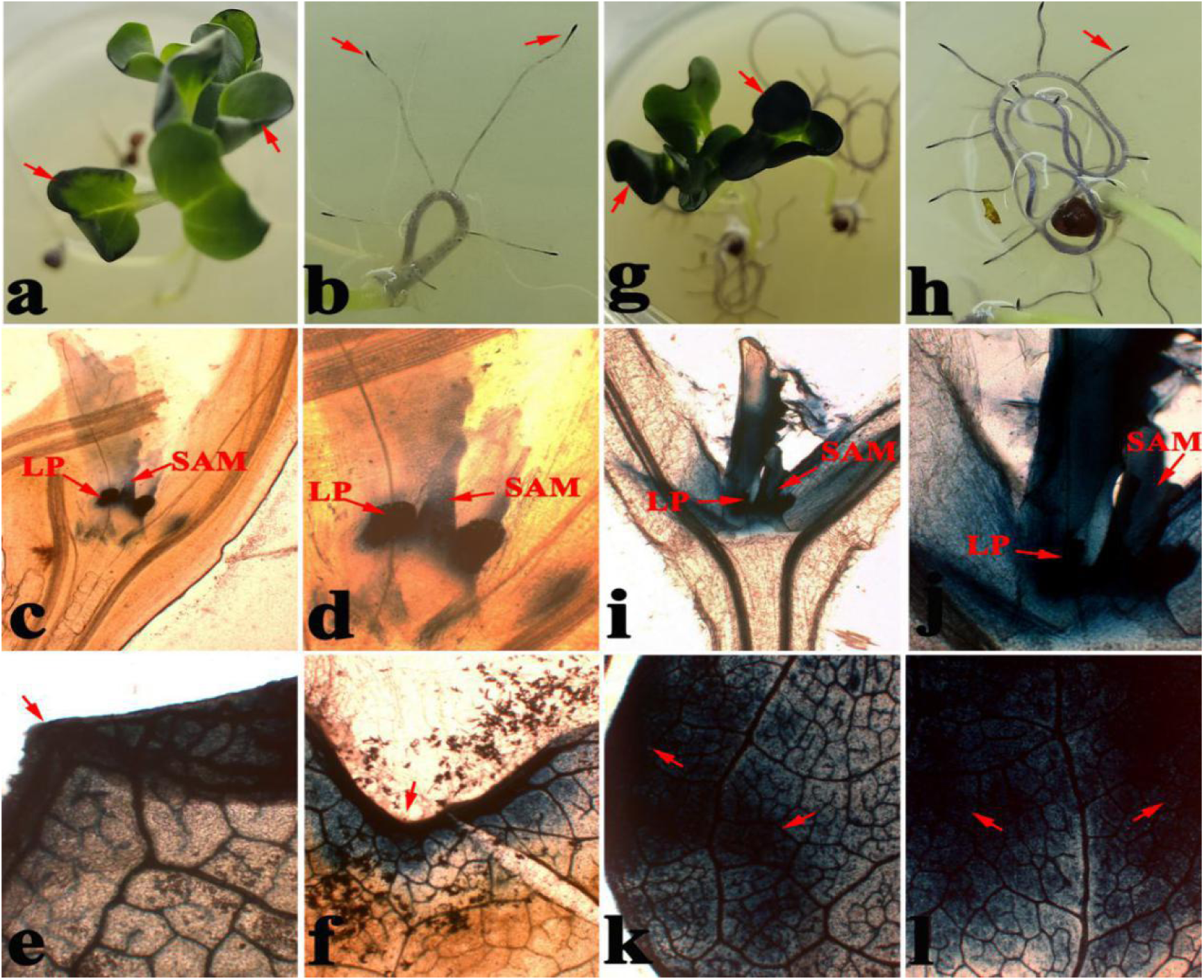
To determine the distribution of O^2.−^ in the leaves, plant tissues were stained with nitroblue tetrazolium (NBT), which is widely used as an indicator of O^2.−^ levels. The O^2.−^ levels in NBT-stained cultivar 16NTS309 seedlings under non-cold (a, b, c, d, e, f) and cold-stress (g, h, i, j, k, l) conditions. SAM: Shoot apical meristems; LP: leaf primordial. Red arrows were indicated super oxide anion (O^2•−^) by dispersion polymerization product of blue spots.

Root apical meristem is responsible for the growth of the primary root[23]. The initiation of lateral roots occurs some distance away from the root apical meristem in the root’s differentiation zone[24]. The mature pericycle cells, once stimulated, differentiate and proliferate to form a lateral root primordium (LRP). The LRP grows through the overlying cell layers of the parent root and eventually breaks through the epidermis and emerges [25]. Under normal conditions, the cluster of cells at the tip of the root (Fig 3b, 4a) and LRP (Fig 4b) expressed greater levels of ROS (O^2.−^). After exposure to cold stress, ROS (O^2.−^) levels increased in the root (Fig 3h, 4c), root tips (Fig 4c), root apical meristems, LRP (Fig 4d, e) and hairy root (Fig 4f) to higher levels than under normal conditions.

**Fig 4.**
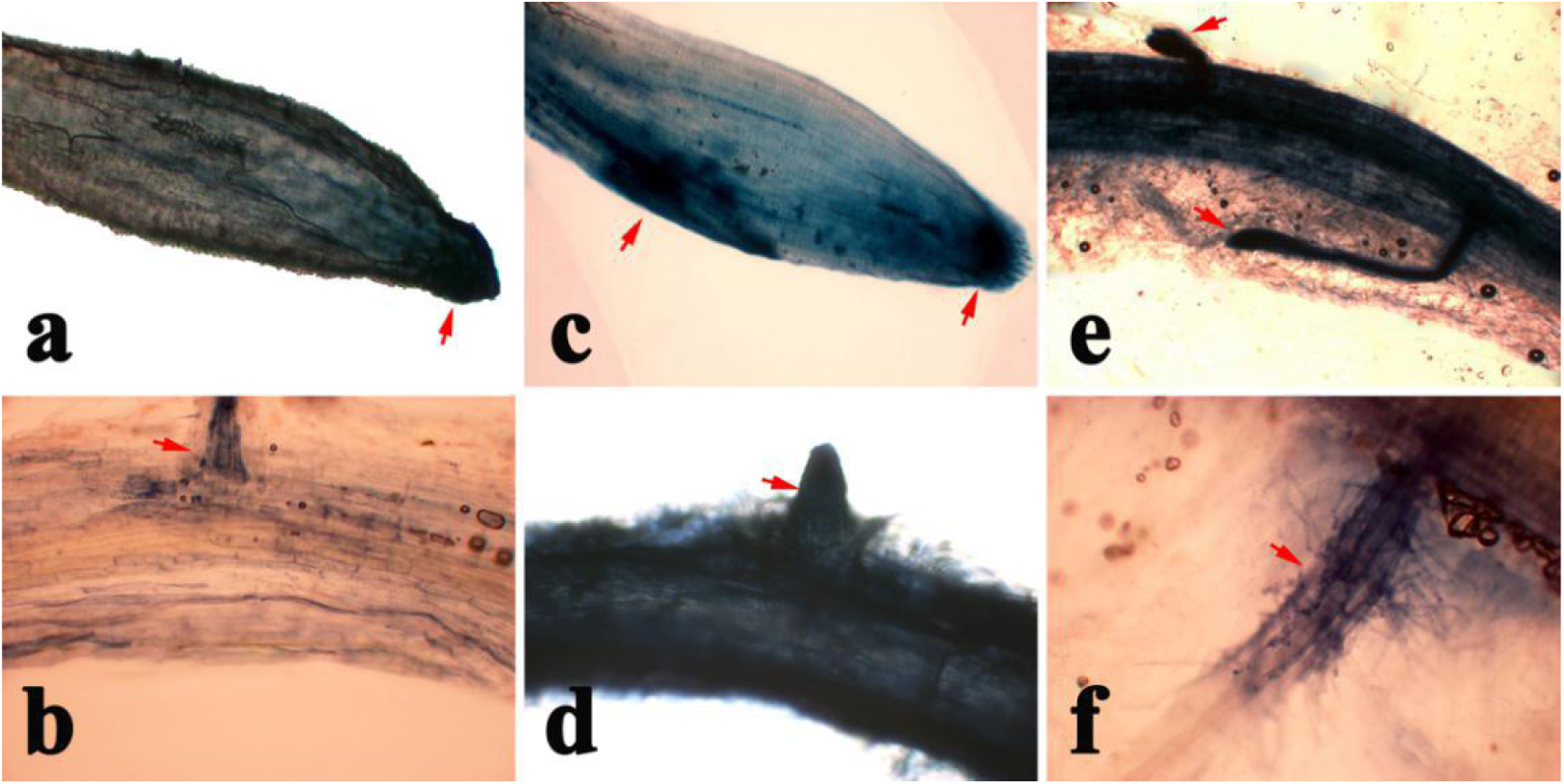
To determine the distribution of O^2.−^ in *Brassica napus* root, we stained plants with nitroblue tetrazolium (NBT), which is widely used as an indicator of O^2.−^ levels. The O^2.−^ levels in NBT-stained cultivar 16NTS309 seedlings under non-cold (a and b) and cold stress (c, d, e f) conditions. lateral root primordium(LRP). Red arrows were indicated super oxide anion (O^2•−^) by dispersion polymerization product of blue spots.

### 2.2 Histological observations of ROS (O^2.−^) in root and leaf cells

The NBT-stained blue sections of roots and leaves from aseptic seedling used for the morphoanatomical analysis were selected for histological observation. Varying degrees of ROS (O^2.−^) accumulation occurred in root and leaf cells under both normal and cold-stress conditions. Under normal conditions, ROS (O^2.−^) were mainly distributed in meristematic cells along leaf edges (Fig 5a) and cambium cells of roots (Fig 6a). The histology of these cells was similar to cells having a strong capacity to proliferate. ROS signals were not detected in other cells.

**Fig 5.**
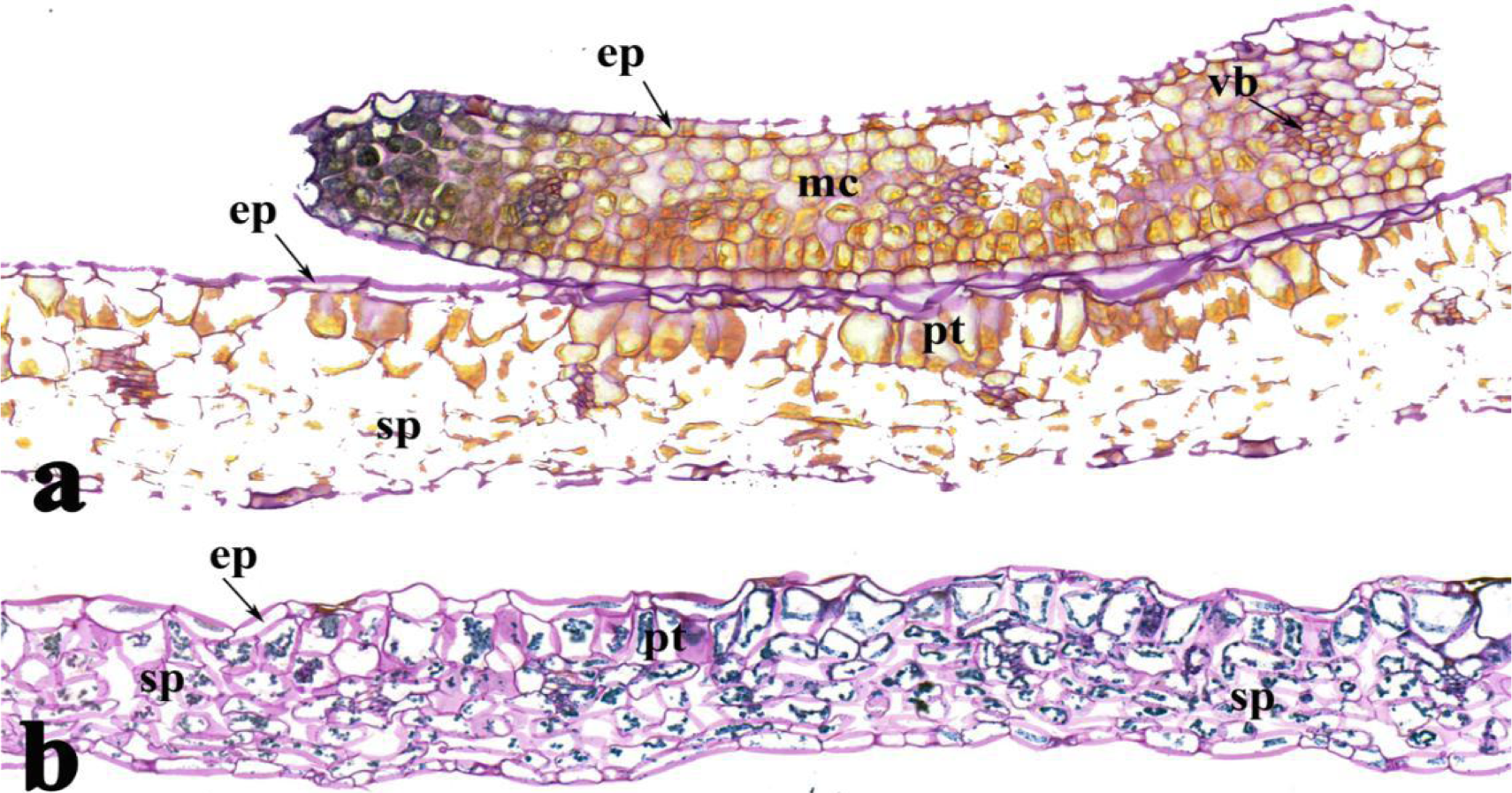
*Brassica napus* cultivar 16NTS309 subjected to non-cold (a) and cold-stress (b) conditions for 48 h. The regions of 16NTS309 leaves samples containing the most blue formazan precipitate were sectioned for morphoanatomical analyses. ep, epidermis; sp, secondary phloem; mc, mesophyll cell; pt, palisade tissue; vb, vascular bundle.

**Fig 6.**
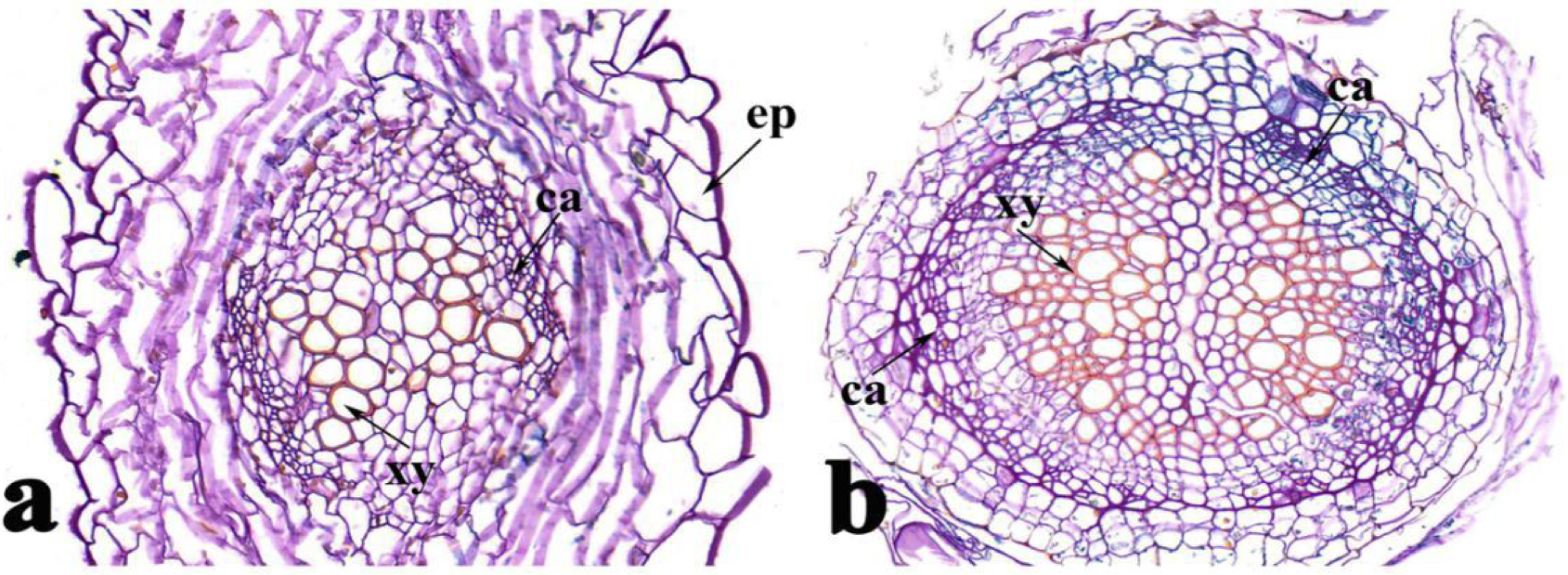
*Brassica napus* cultivar 16NTS309 subjected to non-cold (a) and cold-stress (b) conditions for 48 h. The regions of 16NTS309 roots samples containing the most blue formazan precipitate were sectioned for morphoanatomical analyses. ep, epidermis; ca, cambium; xy, xylem.

After exposure to a cold stress of 4°C, ROS accumulations in cells of roots (Fig 6b), leaves (Fig 5b) were obviously greater than under normal conditions. Leaves are composed of an epidermis, mesophyll and veins. The mesophilic cells near the upper epidermis are cylindrical, forming the palisade tissue, which contains more chloroplasts. The mesophyll cells near the lower epidermis are irregular in shape and loose in arrangement. A greater ROS content accumulated in palisade cells compared with that in other leaf cells, possibly because the latter form spongy tissues containing few chloroplasts.

The root structure, including the epidermis, secondary phloem, secondary xylem, cambium and vascular cylinder, has wide parenchymatous rays. Cambium, which is located between the phloem and xylem in roots, had a large ROS content. LRP initiation is the appearance of closely spaced cell walls in the cambial layer in a perpendicular orientation to the root axis (Fig 6b). The increased cell division frequency and ROS accumulation were clearly seen when compared with cambial cells from the opposite side of the stele. These phenomena confirmed that ROS plays an important role in regulating the balance between cell proliferation and differentiation, by promoting root, stem and leaf growth.

### 2.3 Histological observation of ROS (O^2.−^) in tissues

All plant cells have been traditionally thought of as being totipotent because plant tissues can be induced to form shoots and roots[26-28]. Consequently, we selected the NBT-stained blue sections of calli for histological observation. In vitro, plant tissues having a greater cell-division capacity had greater ROS accumulations under normal conditions, including meristematic nodular tissues, cambium, vascular bundles and root primordium (Fig 7a), while there was little or no ROS accumulation in other cells (Fig 7b). After exposure to cold stress, the ROS distributed in meristematic nodular tissues, cambium, vascular bundles (Fig 7c, d), and other cells increased greatly (Fig 7e). This phenomenon indicated that ROS plays an important role in regulating the balance between cell proliferation and differentiation. ROS signals were detected mainly in cell walls, possibly because ROS have essential functions in broken cell walls and in promoting the formation of different tissues and organs during plant growth.

**Fig 7.**
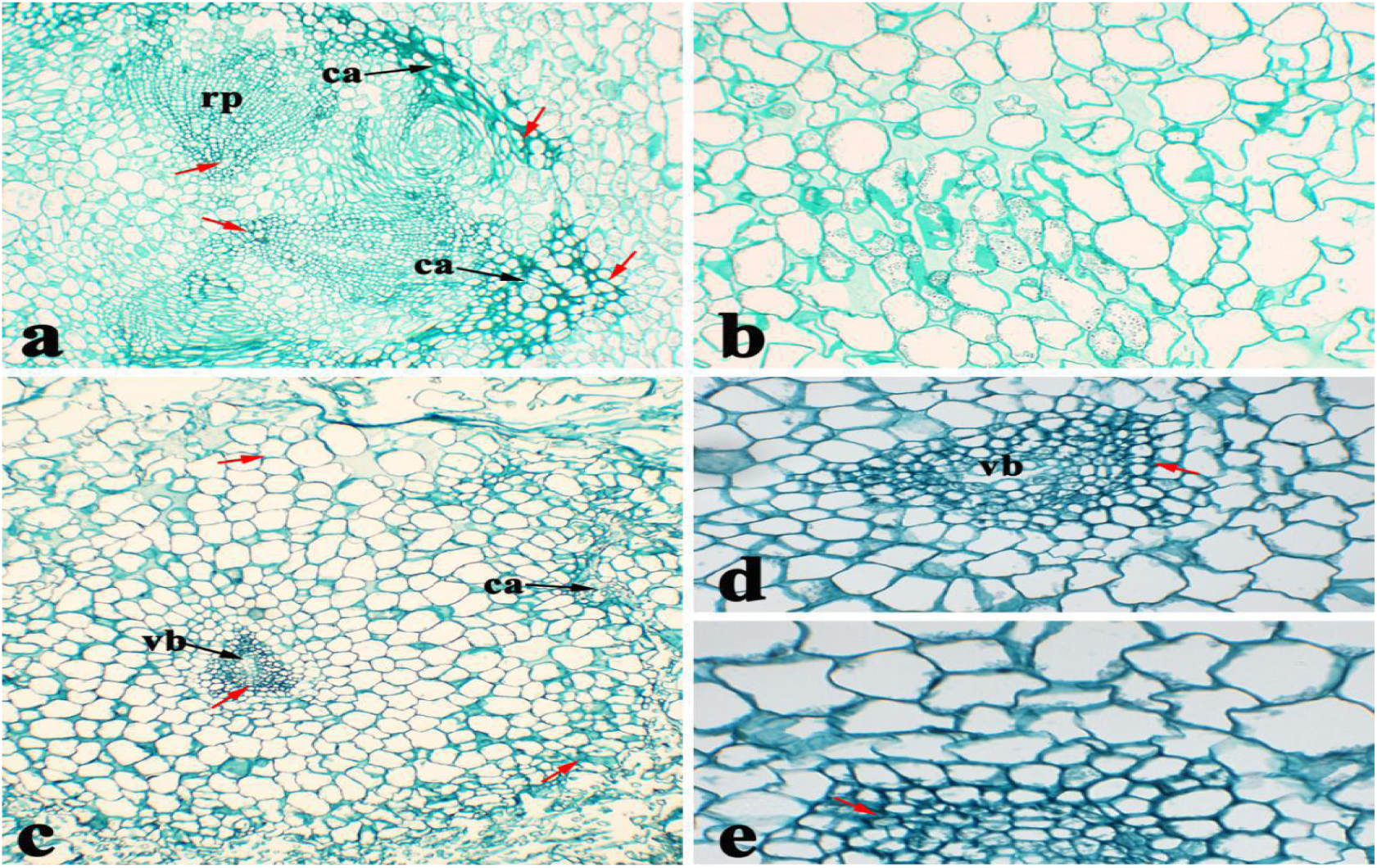
The areas of calli samples having the greatest amount of blue formazan precipitate in *Brassica napus* cultivar 16NTS309 were sectioned for morphoanatomical analyses. ca, cambium; ep, epidermis; vb, vascular bundle; rp, root primordium. Red arrows were indicated super oxide anion (O^2•−^) by dispersion polymerization product of blue spots.

### 2.4 Histochemical staining of onion

ROS is produced intracellularly, but is it also produced extracellularly? To answer this question, we conducted our histochemical staining experiments using onions. After exposure to cold stress, ROS levels increased in onion cells (Fig 8a–e) compared with under normal conditions (Fig 7f). Plasmolysis generally occurs in cells during cold stress. ROS are not only produced in the intercellular space, but are also distributed in the extracellular space or in the cell membrane. ROS staining changed from dark to light (Fig 8). Deep blue-stained areas were seen in each cell, even though ROS had flooded the entire cell. ROS diffused from these dark blue-stained centers to other cells. Thus, we speculated that ROS are first produced outside the cell and then act as signaling molecules to stimulate the production of ROS intracellularly and in other surrounding cells.

**Fig 8.**
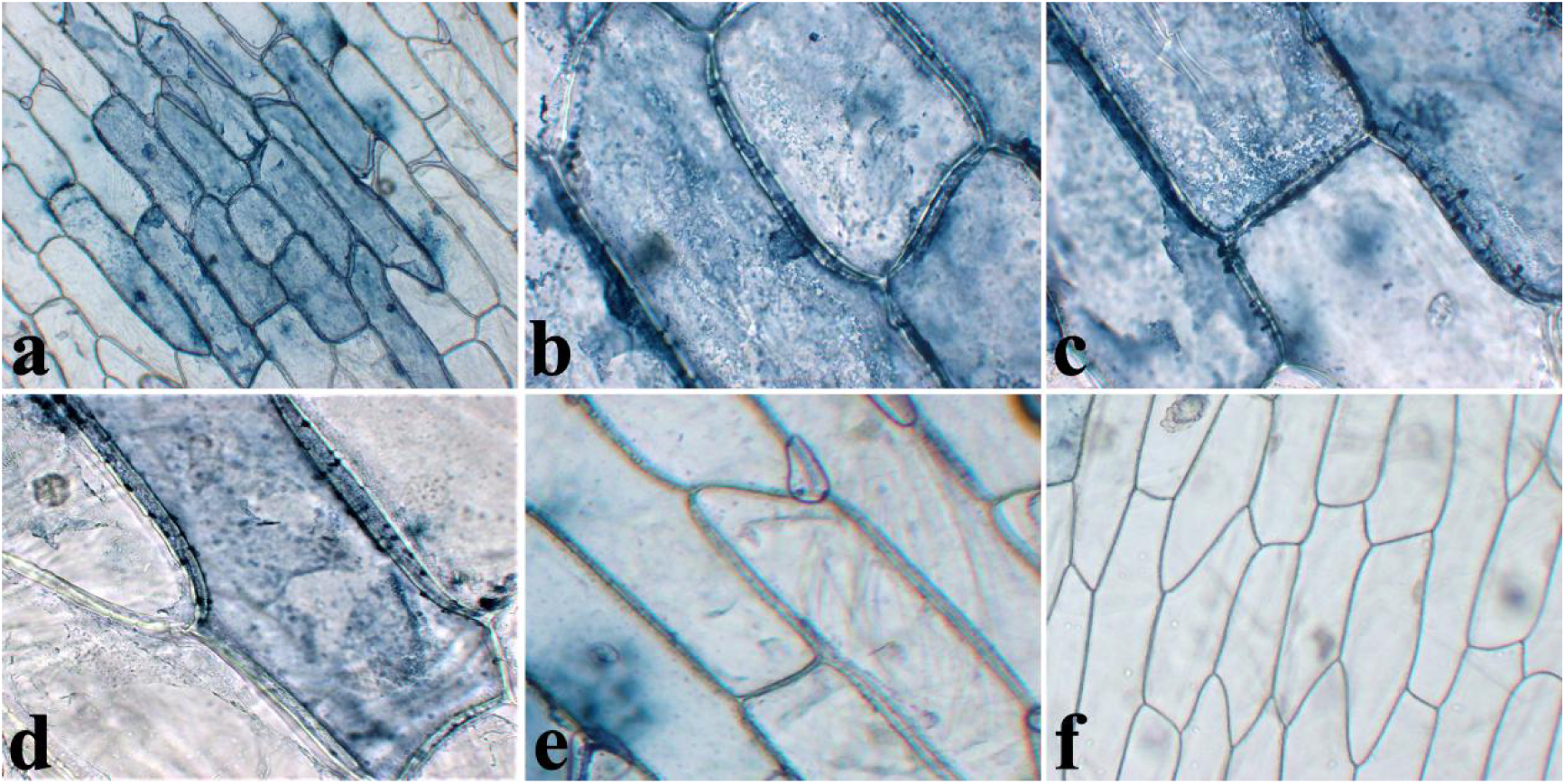
To determine the intracellular and extracellular distribution of O^2.−^, we stained onions with nitroblue tetrazolium (NBT), which is widely used as an indicator of O^2.−^ levels. The onion samples under cold-stress (a–e) and non-cold (f) conditions for 48 h were sectioned for morphoanatomical analyses.

## 3 Discussion

With the shift in our understanding of ROS from toxic chemical species to critical signaling molecules, it is important to elucidate the roles of ROS signaling, which could provide new insights into breeding cold-resistant crops.

In higher plants, organ formation occurs continuously and repetitively from shoot and root apical meristems, which are characterized by high cell division rates[24, 29]. More importantly, is the increase in volume of root, leaf and plant cells. This requires the loosening of the cell wall matrix. However, it has been proposed that ROS regulate the cell proliferation cycle[9]and have direct effects on the stiffness of the plant cell wall [30]. During normal metabolic processes, plant cells produce a variety of ROS, including the O^2.−^, H_2_O_2_ and hydroxyl radicals [31]. Here, under normal conditions, O^2.−^ mainly accumulated in leaf edges (Fig 3a, e, f), SAM and LPs (Fig 3c, d), root tips (Fig 3b, 4a), LRPs (Fig 4b), calli of meristematic nodular tissue, cambia, vascular bundles and root primordia (Fig 6a). These cells are characterized by high division rates. This phenomenon confirmed that ROS plays an important role in regulating the balance between cell proliferation and differentiation, as indicated by the histological staining of tissue sections from plant sterile seedlings and calli of *B. napus* cultivar 16NTS309. The results were verified by assessing the histological staining of onion cells. Our results also confirmed that ROS appeared in the intracellular and extracellular zones, and mainly in the cell wall or membrane, which depends on ROS generation by plasma membrane-localized NADPH oxidases (respiratory burst oxidase homologs), cell wall peroxidases and amine oxidases [32, 33]. This indicated that O^2.−^ preferentially accumulates in the cell wall or membrane zone. More importantly, the results support a dual role for apoplastic ROS in which they have a direct effect on the stiffness of the cell wall, because they cleave cell-wall polysaccharides, and they act as wall loosening agents, thereby either stimulating or restricting cellular extension [34, 35], which promotes root and leaf growth and differentiation [11, 12].

Abiotic stresses, including osmotic, heavy metal ions and dehydration, are important factors that induce somatic embryogenesis in *Arabidopsis* [36]. H_2_O_2_ treatments induce somatic embryogenesis, indicating a role for ROS regulation during this process [37]. Recent evidence indicates a potential promotive effect of oxidative stress on the initiation of somatic embryogenesis [38]. After cold stress, ROS levels increased in leaf (Fig 3g, k, l), root (Fig 3h), LPs and SAM (Fig 3 i, j), root tips, root apical meristem ((Fig 4c), LRPs (Fig 4d, e), hairy roots (Fig 4f), most calli cells(Fig 7c, d, e) and cambia and vascular bundles (Fig 6b). Because of the ubiquitous role of ROS during abiotic stress signaling and development, these results are not surprising. It was also speculated that a moderate low temperature is an important factor for ROS production, after which ROS play roles in promoting cell division and differentiation. Lastly, new shoots and strong root system formed to increase the adaptability to cold stress. A greater ROS content is not always better for plant health, because excessive ROS accumulations from multiple sources result in “ROS bursts” that damage surrounding cells and severely damage cellular structures [39-42].

## Acknowledgement

This study was financially supported by the Agriculture Research System of China (CARS-12); The Agriculture Research System of Gansu Province (GARS-TSZ-1); The Youth Program of Chengdu Normal University (CS19ZC03);

## Author Contribution statement

WL Q and WC S conceived and designed the study. WLQ, W F, L M, P W conducted the experiments. WL Q, WC S analyzed the data.WL Q, JY W, JC W, JJ J contributed reagents, materials, and analysis tools. WLQ wrote the manuscript.We thank Lesley Benyon, PhD, from Liwen Bianji, Edanz Group China (www.liwenbianji.cn/ac), for editing the English text of a draft of this manuscript.

## Conflicts of interest statement

The authors declare no conflict of interest.

